# Reduced Prefrontal PRDM2 Promotes Stress-Induced Alcohol Reinstatement Across Sexes through a dmPFC-Nucleus Accumbens Pathway

**DOI:** 10.64898/2026.03.12.711276

**Authors:** Kanat Chanthongdee, Nicola Murgia, Li Xu, Tetiana Kardash, Sanne Toivainen Eloff, Andrea Coppola, Sofia Vellere, Esi Domi, Estelle Barbier

## Abstract

Stress is a major driver of relapse in alcohol use disorder (AUD), partly through stress-induced dysregulation of prefrontal cortex (PFC) circuits that govern executive control, craving regulation, and adaptive behavioral responses. Such dysfunction may be partly driven by epigenetic regulation of transcriptional programs that shape PFC responses to stress. Here, we investigated the role of the histone methyltransferase PRDM2 in stress-induced relapse to alcohol seeking. Analysis of postmortem human tissue showed that *PRDM2* expression in the PFC was reduced in both men and women with AUD compared with control individuals. To examine the functional significance of this reduction in alcohol-related behaviors, we used viral-mediated knockdown of *Prdm2* in the dorsomedial prefrontal cortex (dmPFC) of male and female rats. *Prdm2* knockdown increased vulnerability to stress-induced reinstatement of alcohol seeking in both sexes, without altering pain sensitivity or being influenced by estrous cycle stage. To determine whether this effect was mediated through specific prefrontal output pathways, we selectively reduced *Prdm2* expression in dmPFC neurons projecting to the nucleus accumbens (NAc). Projection-specific knockdown also increased stress-induced reinstatement of alcohol seeking in male and female rats in a shock intensity-dependent manner. Together, these findings suggest that reduced *PRDM2* expression in the PFC contribute to stress-induced relapse-like behavior and identify the dmPFC-NAc projection as a circuit through which PRDM2 influences alcohol seeking.

## INTRODUCTION

Alcohol use is a major contributor to the global disease burden, leading to substantial mortality and disability worldwide. Relapse remains one of the most persistent challenges in treating AUD, contributing to its chronic, relapsing course (Brady and Sonne, 1999; Sinha et al., 2011; Sliedrecht et al., 2019). Clinical studies consistently identify stress exposure as a powerful trigger for increasing alcohol craving and relapse (Sinha, 2012). In addition, preclinical research shows that stress robustly reinstates alcohol seeking after extinction (Le et al., 1998), suggesting that stress-responsive neural circuits contribute to relapse-like behavior. Converging evidence implicates dysfunction of the PFC in stress-induced alcohol relapse, particularly through impaired top-down regulation of subcortical reward and stress-related circuits (Barbier et al., 2017; Blaine et al., 2017; Lu and Richardson, 2014; Seo et al., 2013). However, the molecular mechanisms within defined PFC circuits that confer vulnerability to stress-induced alcohol relapse remain poorly understood, limiting the development of targeted therapeutic strategies.

Substantial evidence indicates that epigenetic regulators play a central role in shaping stress responsivity and alcohol-related behaviors by modulating transcriptional programs within corticolimbic networks (Barbier et al., 2017). Both stress and alcohol exposure can induce persistent epigenetic modifications that alter gene expression without changing the DNA sequence, leading to long-lasting adaptations in neural circuit function (Barbier et al., 2015; Bohnsack et al., 2022). Among these regulators, the histone methyltransferase PRDM2 has emerged as an important regulator of behavioral control in the medial prefrontal cortex (mPFC). For instance, chronic alcohol exposure reduces *Prdm2* expression in the dmPFC. Moreover, viral vector-mediated knockdown of *Prdm2* is sufficient to enhance alcohol-seeking behavior, indicating that PRDM2 functionally regulates alcohol seeking. PRDM2 also regulates the expression of genes involved in synaptic plasticity, providing a potential mechanistic link between transcriptional regulation and alcohol-related behaviors (Barbier et al., 2017). These findings support a role for PRDM2 as a key epigenetic mediator linking transcriptional regulation to maladaptive behavioral responses to alcohol and stress. However, several critical questions remain. In particular, it remains unclear how PRDM2 integrates stress-related signals within specific prefrontal circuits and whether its effects are conserved across sexes and species, leaving important aspects of its role in stress-induced reinstatement unresolved.

Notably, *PRDM2* expression in the human PFC has not been examined in AUD, and its behavioral effects have thus far been studied exclusively in males. Addressing these gaps is important, both for establishing the clinical relevance of PRDM2 in AUD and for determining whether its role generalizes across sexes, particularly given well-established sex differences in stress responsivity and alcohol-related behaviors (Peltier et al., 2019). Furthermore, prior studies have not determined whether PRDM2 influences alcohol relapse through global modulation of dmPFC activity or through specific downstream projections. Although dmPFC activity has been implicated in stress-induced reinstatement of alcohol seeking, the specific prefrontal output pathways mediating this effect remain poorly defined (Barbier et al., 2017). The dmPFC projection to the nucleus accumbens (NAc) is a strong candidate, as this pathway plays a critical role in stress-induced cocaine relapse and the integration of executive control with motivational drive (McFarland et al., 2004; Yang et al., 2021). Given the established role of the dmPFC in alcohol relapse, the dmPFC→NAc pathway emerges as a plausible circuit through which prefrontal activity may regulate stress-induced alcohol reinstatement.

To address these questions, we combined postmortem human analyses with circuit-specific manipulations in male and female rodent models of stress-induced reinstatement of alcohol seeking. We first demonstrate that *PRDM2* expression is reduced in the PFC of both men and women, establishing translational relevance and sex convergence at the molecular level. We then show that *Prdm2* knockdown in the dmPFC enhances stress-induced reinstatement of alcohol seeking in both male and female rats, extending prior male-only findings. Finally, using projection-specific knockdown strategies, we identify the dmPFC→NAc pathway as a critical circuit through which PRDM2 promotes stress-induced alcohol relapse behavior. Together, these findings identify PRDM2 as an epigenetic regulator of stress-induced alcohol relapse acting within a defined prefrontal-striatal circuit across sexes.

## MATERIAL AND METHOD

### Postmortem human samples

Adult post-mortem PFC of humans diagnosed with AUD and their controls, including both men and women, were obtained from the New South Wales Brain Bank. All procedures with post-mortem human samples were conducted in accordance with ethical guidelines and approved by The Swedish Ethical Review Authority (Dnr: 2017/407-31).

### ANIMALS

Adult male (250–300 g) and female (150–200 g) Wistar rats (Charles River, Germany) were housed under a reverse light cycle with *ad libitum* access to food and water. All experimental procedures were conducted in accordance with guidelines established by the Swedish National Committee for animal research and were approved by the Local Ethics Committee for Animal Care and Use at Linköping University.

### BEHAVIORAL TESTING

#### Alcohol self-administration

Rats were trained to self-administer 20% (v/v) alcohol in water following previously established protocols (Augier et al., 2014). Rats were first trained on a fixed-ratio 1 (FR1) schedule during 30 min sessions. Each lever press delivered 100 µL of 20% alcohol into the drinking cup receptacle and triggered a 5s time-out period, during which additional presses did not result in alcohol delivery. After stable baseline responding was achieved, the schedule was increased to fixed-ratio 2 (FR2). After stabilizing self-administration under the FR2 schedule, rats were bilaterally infused in the dmPFC with either a PRDM2-targeting viral vector or a scrambled control vector (viruses details are provided in the surgeries section). A second experimental group received a Cre-dependent PRDM2-targeting vector or Cre-dependent scrambled vector in the dmPFC together with retro-Cre-EGFP virus into the NAc (virus details are provided in the surgeries section), enabling pathway-specific manipulation of dmPFC→NAc projections. Alcohol self-administration at FR2 was then resumed 1 month after surgery to allow for viral vector expression (Fig.1B).

#### Stress-induced reinstatement

Stress-induced reinstatement was performed as described previously (Barbier et al., 2017). Briefly, extinction sessions started once self-administration rates were stable. During extinction, alcohol was no longer available, and operant responses had no programmed consequences. Stress-induced reinstatement testing started after operant responding was extinguished (mean interval around 10 active lever presses per 30 min session). Rats then underwent three reinstatement testing, each following 15-minute exposure to randomized electric footshocks at 0.4, 0.6 or 0.8 mA. Each test was separated by 3-5 days of extinction training (Fig.1B).

#### Foot shock sensitivity

Foot shock sensitivity thresholds were assessed in Med Associates operant chambers as previously described (Domi et al., 2021). Rats received 0.5 s foot shocks in 0.1 mA increments every 30s, beginning at 0.1 mA, and an observer blinded to the experimental groups recorded the withdrawal of one, two, or all four paws.

#### Surgeries

Rats received microinjections of AAV vectors using stereotaxic surgery under anesthesia induced with isoflurane (3–5 %) and maintained at 1.5–2.5 % for the duration of the procedure. All AAV vectors were manufactured from Neuroscience Center Zurich, University of Zurich and ETH Zurich, Zurich, Switzerland. Targets for AAV injections were used according to Paxinos Atlas. Coordinates relative to bregma; dmPFC: anterior-posterior: +2.76 mm, medial-lateral: ± 0.6 mm, dorsal-ventral -3.5 mm; NAc: anterior-posterior: 1.8 mm, medial-lateral: ±1.4 mm, dorsal-ventral -7 mm; (Paxinos and Watson, 2006) (Fig. S1).

##### Prdm2 KD in the dmPFC neurons

Rats received bilateral microinjections of an AAV containing shRNA targeting *Prdm2* (AAV5-hSyn-shRNA-Prdm2-mScarlet; original titer 7.7 × 10^12^ VG/ml in sterile 1X PBS at 1:100 dilution titer) or a scrambled control (AAV5-hSyn-shRNA-mScarlet; titer 1.0 × 10^13^ GC/ml in sterile 1X PBS at 1:100 dilution titer) into the dmPFC (0.75 μL per injection; rate: 0.1 μL/min).

##### Prdm2 KD in the dmPFC→NAc

Rats received microinjections of two AAV vectors for intersectional gene manipulation approach. Rats received bilateral injections of an AAV containing Cre-dependent shRNA targeting *Prdm2* (AAV5-hSyn-mScarlet-shPrdm2-DIO; original titer 1.1 × 10^13^ VG/mL in sterile 1X PBS at 1:10 dilution titer) or a Cre-dependent scrambled control (AAV5-hSyn-mScarlet-scr-DIO; titer 7.0 × 10^12^ GC/ml in sterile 1X PBS at 1:10 dilution titer) into the dmPFC (0.75 μL per injection; rate: 0.1 μL/min). In the same surgery, rats received bilateral injections of a retrogradely transported AAV encoding Cre-EGFP (ssAAV-retro/2-hSyn1-EGFP_iCre-WPRE-hGHp(A)); titer 7.8 × 10^12^ GC/ml) into the NAc (0.75 μL per injection; rate: 0.1 μL/min).

#### Estrous cycle assessment

To assess potential impacts from hormonal variations across the estrous cycle on stress-induced reinstatement of alcohol seeking, vaginal cellular samples were obtained on the day of stress-induced reinstatement testing at least 1 h following the test session as previously described (Toivainen et al., 2024). A moist cotton swab was rolled against vaginal wall to retrieve cellular samples. The swab was then spread onto a microscope slide and air-dried for subsequent cytological examination. Slides were fixed in 99.5 % ethanol for 10 min, stained for 1 min with a 0.1 % (v/v) Cresyl Violet working solution prepared from a 1 % Cresyl Violet acetate stock (Sigma-Aldrich, C5042-10G), and rinsed with tap water. Vaginal cytology was viewed at x10 and x20 magnification under bright field illumination (ZEISS Primostar 3 microscope) using the corresponding definitions to determine the stage of estrous cycle (Byers et al., 2012). When females are in proestrus, clusters of rounded, well-defined nucleated epithelial cells are mostly observed. This stage is associated with the highest level of plasma progesterone and estrogen (Becker et al., 2005). When the stage advances to estrus, cornified epithelial cells are primarily present. Metestrus, which is the shortest stage, is characterized when cornified epithelial cells are alongside small leukocytes. Diestrus, which is the longest stage, is characterized by the presence of leukocytes in the majority with a few epithelial cells. Subjects were grouped as either proestrus (i.e., high reproductive hormone level) or non-proestrus (i.e., low reproductive hormone level) for subsequent analyses.

#### Quantitative real-time PCR

DNA and RNA were extracted from brain human samples (Fig. 1A) using AllPrep DNA/RNA Mini Kit (Qiagen, MD, USA) and stored at –80 °C. RNA was reverse transcribed into cDNA using High-capacity cDNA Reverse Transcription Kit (Thermo Fisher Scientific, Vilnius, Lithuania). Inventoried TaqMan Gene expression assay probes (*PRDM2*: Hs00202013_m1 and *ACTB*: Hs01060665_g1; Life Technologies, CA, USA) and TaqManTM Fast Universal PCR Master Mix (Thermo Fisher Scientific, Vilnius, Lithuania) were used to assess gene expression on a QuantStudio 7 Flex Real-time PCR system (Life Technologies, CA, USA). *PRDM2* gene expression was quantified with respect to beta actin (*ACTB*) using 2^− ΔΔCT^ analysis.

**Figure 1:**
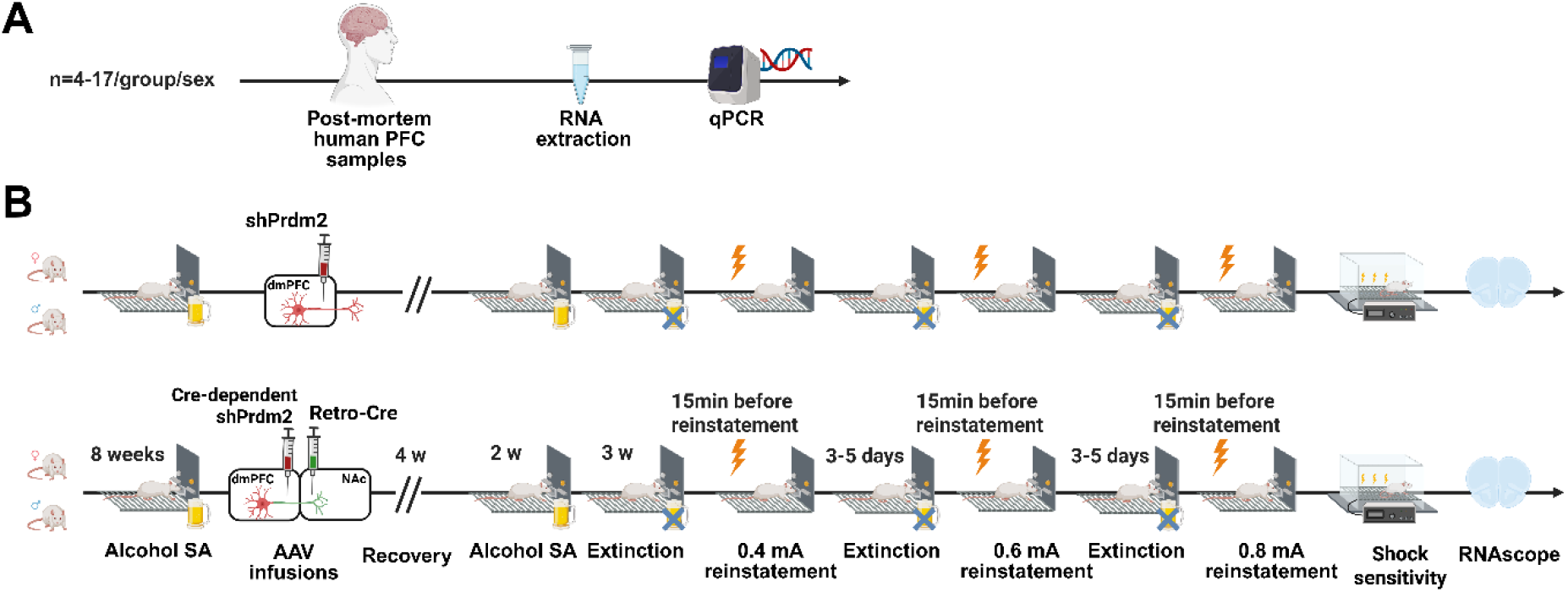
Experimental timeline. (A) Postmortem human prefrontal cortex (PFC) tissue was collected and analyzed to measure *PRDM2* expression using quantitative real-time PCR. (B) Experimental design in rats. Male and female rats were trained for alcohol self-administration followed by stereotaxic viral injections targeting the dorsomedial prefrontal cortex (dmPFC) or dmPFC neurons projecting to the nucleus accumbens (NAc) to induce *Prdm2* knockdown (KD) or scrambled control expression. After recovery, rats underwent alcohol self-administration training (fixed-ratio schedule) followed by extinction sessions. Stress-induced reinstatement of alcohol seeking was then tested using intermittent footshock stress at increasing intensities (0.4, 0.6, and 0.8 mA) across separate test sessions. Behavioral responding during reinstatement was compared with the preceding extinction baseline. After completion of behavioral testing, brain tissue was collected for verification of viral expression and molecular analyses.

#### RNAscope^®^ in situ hybridization

To assess *Prdm2* expression in the dmPFC, brains were flash frozen in pre-chilled isopentane and sectioned using a cryostat to achieve 16 µm thickness. The sections were processed following RNAscope® Multiplex Fluorescent v2 assay according to the manufacturer’s protocol. Briefly, slides were first dehydrated using a series of ethanol and treated with protease. Transcripts on the slides were hybridized using *Prdm2* probe (ACDBio cat# 457861). Slides were then serially incubated in signal amplification molecules, channel-specific HRP, Tyramide Signal Amplification fluorophores (Cy5 channel for *Prdm2* transcript), and HRP blocker. Lastly, slides were incubated in DAPI, coverslipped, and stored at 4 °C at least overnight before imaging. Photomicrographs were acquired with Zeiss LSM 800 upright confocal microscope at x20 objective lens. One photomicrograph from the dmPFC in each hemisphere was acquired using identical laser and pinhole settings. Cells and subcellular puncta were counted using an automated quantification software (QuPath version 0.5.1) that was set with consistent thresholding levels. Cells were identified using DAPI channel (detection threshold=2000) and subcellular puncta were detected using Cy5 channel (detection threshold=13200, 0.5-8 pixels in diameter, located within 3 µm expansion from the DAPI outlines). Each punctum was assumed to represent one *Prdm2* transcript. The mean *Prdm2* expression in the dmPFC were calculated and reported as transcripts/cells.

### Statistical analysis

All data were analyzed using GraphPad Prism version 10 (GraphPad Software, San Diego, CA) and STATISTICA (StatSoft 13.0 RRID: SCR_014213). Animals lacking confirmed viral expression in the target region were excluded from analysis. From the dmPFC cohort (initial N=48 males, N=64 females), N=5 males and N=3 females were excluded, leaving N=43 males and N=61 females for analysis. From the dmPFC→NAc cohort (initial N=48 males, N=48 females), N=6 males and N=4 females were excluded, leaving N=42 males and N=44 females. One female rat from the dmPFC cohort was excluded from the 0.6 mA reinstatement test due to injury sustained during testing. Homogeneity of variance was assessed using the Levene test. When data violated ANOVA assumption, statistical analysis was performed using aligned rank transform (ART) ANOVA using ART tool package in R (Kay, 2025; Wobbrock, 2011). All analyses were conducted in R (R Core Team) using RStudio. When no violation of assumption was observed, parametric ANOVA was used, with factors for each analysis indicated in the result section. Post hoc comparisons were performed using Newman-Keuls test. Statistical outliers were identified and excluded using Grubbs’ test (α=0.05), applied independently to each experimental group, with a maximum of one outlier excluded per group (total N excluded by Grubbs = 8). The accepted level of significance for all tests was *p* < 0.05. Stars indicate significance relative to the scrambled or control groups (* *p* < 0.05, ** *p* < 0.01, *** *p* < 0.001), whereas hash symbols indicate significance for within-group comparisons (*# p* < 0.05, *## p* < 0.01, *### p* < 0.001).

## RESULTS

### Reduced *PRDM2* Expression in the Prefrontal Cortex of Men and Women with Alcohol Use Disorder

We obtained postmortem PFC samples from individuals diagnosed with AUD and age-matched controls to measure *PRDM2* expression (Fig. 1A). Demographic characteristics of the study participants are summarized in Table 1. Using quantitative real-time PCR, we found that both men and women with AUD had approximately 30 % lower *PRDM2* expression in the PFC compared with sex-matched control individuals (Fig. 2). Two-way ANOVA indicated a significant main effect of group (AUD vs. controls; F_(1,35)_=14.24; p<0.01) and a significant main effect of Sex (Male vs. Female; F_(1,35)_=22.92; p<0.01). No significant interaction effect was observed (Group X Sex; F_(1,35)_=2.70; p=0.11). These findings indicate that, while women exhibit lower overall *PRDM2* expression than men, AUD is associated with reduction in *PRDM2* expression in both men and women.

**Table 1:** Demographic characteristics of post-mortem prefrontal cortex samples. Summary of sex distribution, age, post-mortem interval, and brain pH for control subjects (N=18) and individuals with alcohol use disorder (N=21). Values are reported as counts and percentages for categorical variables, and as mean ± SD for continuous variables.

| Characteristics | Control<br>(N=18) |  | Alcohol use disorder<br>(N=21) |  |
| --- | --- | --- | --- | --- |
|  | N | % | N | % |
| Males | 14 | 78,2 | 17 | 81,0 |
| Females | 4 | 22,2 | 4 | 19,0 |
|  | Mean | SD | Mean | SD |
| Age (years) | 54,9 | 10,2 | 58,7 | 10,8 |
| Post-mortem interval (hours) | 32,0 | 11,4 | 38,1 | 15,6 |
| Brain pH | 6,6 | 0,3 | 6,5 | 0,3 |

**Figure 2:**
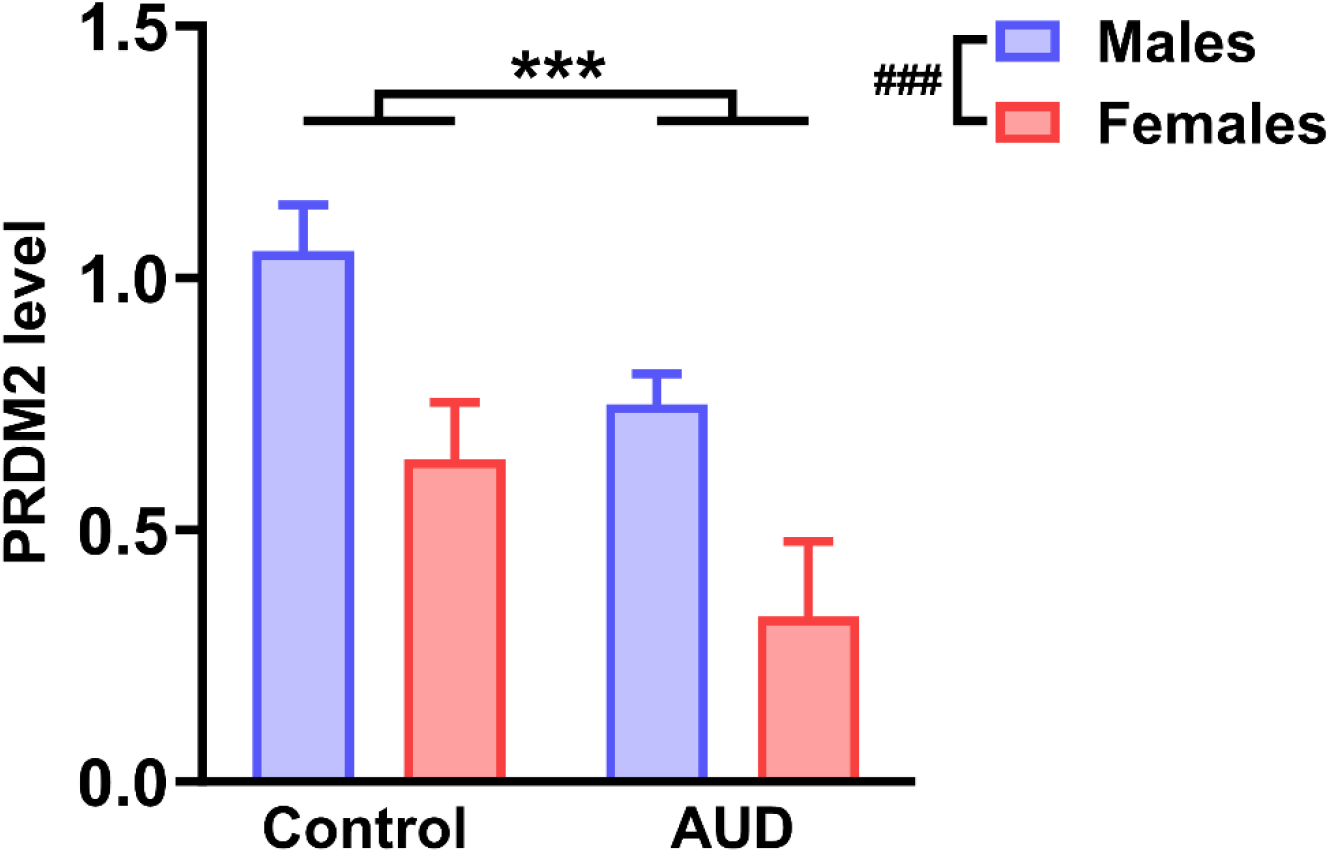
*PRDM2* expression in the medial prefrontal cortex in control and alcohol use disorder (AUD) subjects. Relative PRDM2 levels in the prefrontal cortex of male (blue) and female (red) control subjects and individuals with alcohol use disorder (AUD). *PRDM2* expression was significantly reduced in AUD compared with controls (*** *p* < 0.001). In addition, a significant sex effect was observed, with females showing lower PRDM2 levels compared with males (### *p* < 0.001). Bars represent mean ± SEM.

### *Prdm2* KD in the dmPFC promotes stress-induced reinstatement of alcohol seeking in rats

To determine whether the previously reported role of *PRDM2* in stress-induced reinstatement extends across sexes, we examined the effect of *Prmd2* KD in the dmPFC of both male and female rats on stress-induced reinstatement of alcohol seeking (Fig. 1B). Viral expression was restricted to the dmPFC (Fig. 3A) and resulted in ∼ 30 % downregulation of *Prdm2* expression in the dmPFC compared with scrambled controls (one-way ANOVA: scrambled vs. KD; main effect of group; F_(1,18)_=5.96; p<0.05; Fig. 3B).

**Figure 3:**
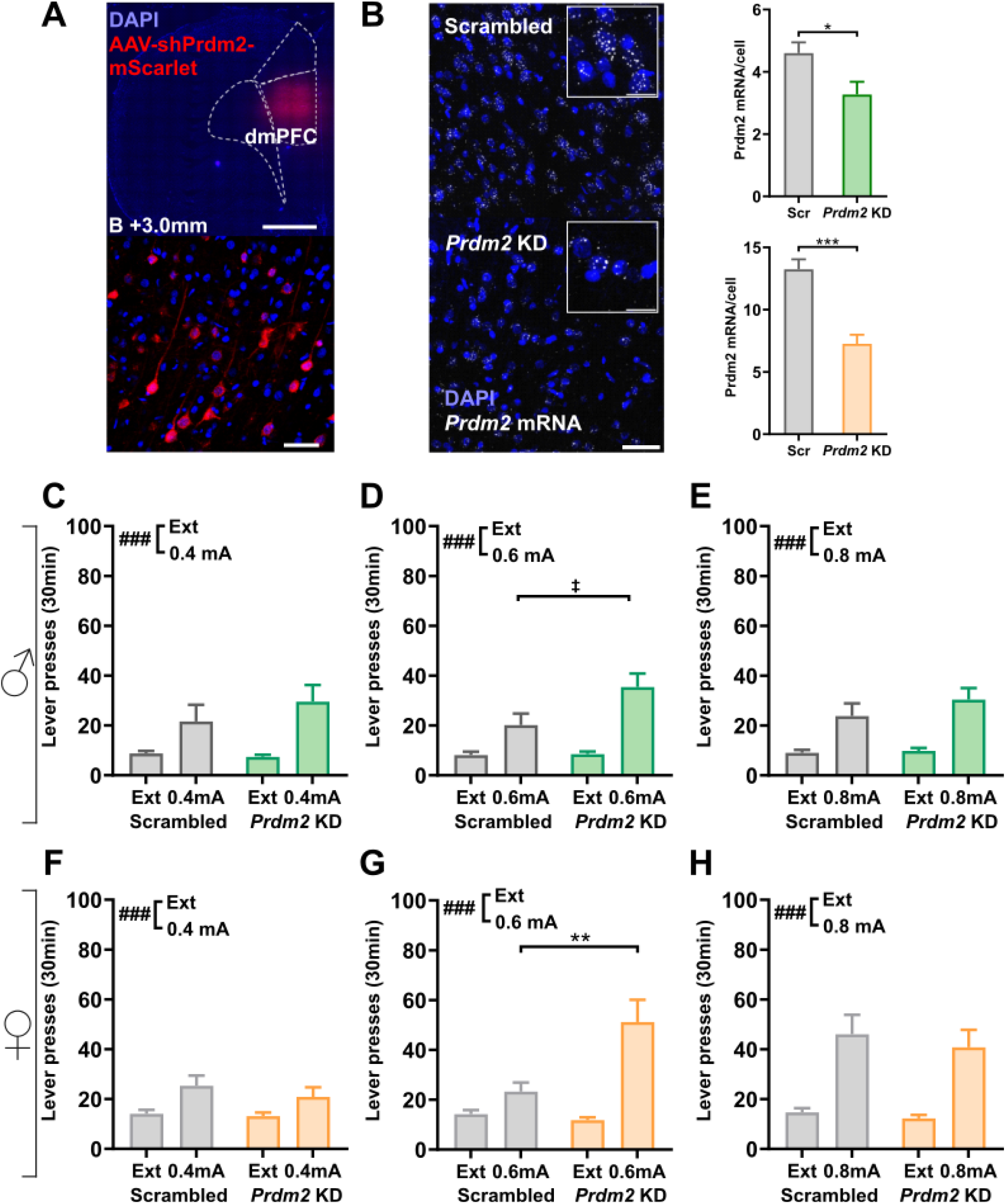
Knockdown of *Prdm2* in the dmPFC enhances stress-induced reinstatement of alcohol seeking at intermediate shock intensity in both male and female rats. (A) Representative images of AAV5-hSyn-shRNA-Prdm2-mScarlet targeting the dmPFC. Top: low-magnification coronal section showing viral expression (mScarlet, red) confined to the dmPFC (dashed outline; Bregma +3.0 mm). Scale bar: 1000 µm. Bottom: higher magnification image of AAV5-hSyn-shRNA-Prdm2-mScarlet expressing neurons (red) with DAPI nuclear counterstain (blue). Scale bar: 50 µm. (B) Validation of *Prdm2* knockdown (KD) in the dmPFC using RNAscope® in situ hybridization. Representative images from scrambled control and *Prdm2* KD animals showing *Prdm2* mRNA puncta (white, Cy5 channel) and DAPI nuclear counterstain (blue). Insets show higher-magnification views of individual cells. Scale bars: 50 µm (main images), 20 µm (insets). Quantification of *Prdm2* mRNA puncta per cell in males (top) and females (bottom) demonstrates significantly reduced *Prdm2* expression in KD animals compared to scrambled controls. Data are presented as mean ± SEM. * *p* < 0.05, *** *p* < 0.001. (C–E) Stress-induced reinstatement of alcohol seeking in male rats following intermittent footshock at 0.4 mA (C), 0.6 mA (D), and 0.8 mA (E). Stress exposure increased lever pressing relative to extinction conditions across shock intensities, with *Prdm2* KD significantly enhancing reinstatement responding at 0.6 mA. (F–H) Stress-induced reinstatement of alcohol seeking in female rats tested at 0.4 mA (F), 0.6 mA (G), and 0.8 mA (H). Similar to males, *Prdm2* KD significantly increased reinstatement responding at 0.6 mA but not at the lower or higher shock intensities. Bars represent mean ± SEM lever presses during the 30-min session. ‡ *p* = 0.05 (compared to scrambled control), ** *p* < 0.01 (compared to scrambled control); ### *p* < 0.001 main effect of stress compared with extinction.

All rats underwent footshock-induced reinstatement testing at 0.4, 0.6, and 0.8 mA shock intensities (males: Fig. 3C–E; females: Fig. 3F–H; Fig. S2). In males, stress exposure significantly reinstated alcohol seeking across shock intensities and replicated our previous findings that *Prdm2* KD enhances stress-induced reinstatement at 0.6 mA. At 0.4 mA, repeated-measures ANOVA showed a significant main effect of stress (F_(1,36)_=13.42, p=0.0008), but no main effect of group (F_(1,36)_=0.46, p=0.50) and no Stress × Group interaction (F_(1,36)_=0.97, p=0.33; Fig. 3C). At 0.6 mA, stress robustly reinstated alcohol seeking (F_(1,36)_=28.09, p<0.001). A significant main effect of group was observed (F_(1,36)_=4.14, p=0.049), and there was a trend toward a Stress × Group interaction (F_(1,36)_=4.08, p=0.051; Fig. 3D). Post hoc comparisons indicated that *Prdm2* KD rats exhibited greater reinstatement responding compared with scrambled controls (p<0.001). At 0.8 mA, stress significantly reinstated alcohol seeking (F_(1,36)_=26.61, p<0.001), but no group effect (F_(1,36)_=1.03, p=0.32) and no Stress × Group interaction (F_(1,36)_=0.74, p=0.40; Fig. 3E).

A comparable pattern was observed in female rats (Fig. 3F-H). At 0.4 mA, repeated-measures ANOVA showed a significant main effect of stress (baseline extinction vs. reinstatement; F_(1,52)_=12.16, p=0.001), indicating increased responding during reinstatement testing. However, there was no main effect of group (scrambled vs. *Prdm2* KD; F_(1,52)_=0.66, p=0.42) and no Stress × Group interaction (F_(1,52)_=0.41, p=0.52; Fig. 3F). At 0.6 mA, because the assumption of homogeneity of variance was violated (Levene’s test: p=0.0005), data were analyzed using an aligned rank transform (ART) ANOVA. This analysis showed a significant main effect of group (F_(1,51)_=9.13, p=0.004), a significant main effect of stress (F_(1,51)_=34.16, p=3.58×10^−7^), and a significant Group × stress interaction (F_(1,51)_=14.41, p=0.0004; Fig. 3G). Post hoc analysis indicated that *Prdm2* KD females showed significantly greater reinstatement responding than scrambled controls (P=0.004). At 0.8 mA, repeated-measures ANOVA revealed a significant main effect of stress (F_(1,53)_=32.40, p<0.0001), but no main effect of group (F_(1,53)_=0.48, p=0.49) and no Stress × Group interaction (F_(1,53)_=0.07, p=0.79; Fig. 3H). In line with our prior findings in males, *Prdm2* KD did not affect pain sensitivity to footshock in females (repeated-measures ANOVA: no main effect of Group, F_(1,21)_=0.29, p=0.59, and no Group × Intensity interaction, F_(2,42)_=0.33,p=0.72; Fig. 4A), indicating that the differences in reinstatement between groups does not result from altered pain thresholds.

**Figure 4:**
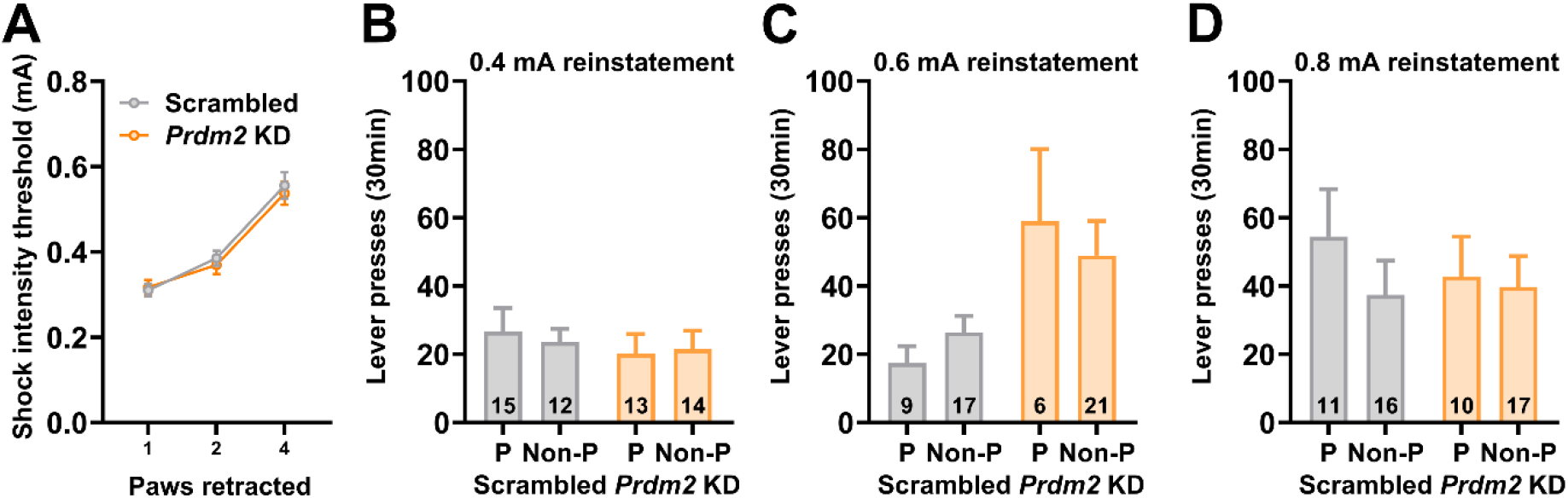
*Prdm2* knockdown does not alter pain sensitivity in female rats and reinstatement effects are independent of estrous cycle stage. (A) Footshock sensitivity thresholds measured as the current intensity required to elicit paw withdrawal responses (1, 2, or 4 paws retracted) in scrambled and *Prdm2* KD female rats. No differences in shock sensitivity were observed between groups, indicating that altered reinstatement behavior is not due to changes in pain perception. (B–D) Effect of estrous cycle stage on stress-induced reinstatement of alcohol seeking following intermittent footshock at 0.4 mA (B), 0.6 mA (C), and 0.8 mA (D). Lever presses during the 30-min reinstatement test are shown for female rats tested during proestrus (P) and non-proestrus (Non-P) stages of the cycle. Estrous stage did not influence reinstatement responding and did not interact with group at any shock intensity, including 0.6 mA, where *Prdm2* KD enhanced reinstatement. Bars represent mean ± SEM. Numbers within bars indicate sample sizes.

Since cycling reproductive hormones can affect behavioral responses to stress in females (Pestana et al., 2023), estrous cycle was also assessed after each reinstatement testing (Fig. 4B-D). We did not find any main effect of the estrous stage or interaction between group and estrous stage at any reinstatement testing, including 0.6 mA testing where we observed the effect of *Prdm2* KD (two-way ANOVA main effect of Estrous stage: Proestrus vs. Non-proestrus: F_(1,49)_=0.00, p=0.96, with no Group × Estrous stage interaction, F_(1,49)_=0.72, p=0.40; Fig. 4C). This suggests that fluctuations in reproductive hormones do not influence the reinstatement-promoting effect of *Prdm2* KD.

Together, these findings demonstrate that reduced *Prdm2* expression in the dmPFC enhances stress-induced reinstatement of alcohol seeking in both male and female rats, identifying PRDM2 as a regulator of stress-induced relapse behavior across sexes.

### PRDM2 in dmPFC→NAc Neurons Regulates Stress-Induced Alcohol Reinstatement Across Sexes

While our results demonstrate that PRDM2 in the dmPFC regulates stress-induced reinstatement across sexes, it remains unclear how PRDM2 integrates stress-related signals within specific prefrontal circuits to promote relapse-like behavior. The dmPFC→NAc projection is a strong candidate, as this pathway plays a central role in stress-induced reinstatement to cocaine seeking and motivational control (McFarland et al., 2004; Yang et al., 2021). We therefore tested whether PRDM2 within dmPFC neurons projecting to the NAc regulates stress-induced reinstatement of alcohol seeking by selectively reducing *Prdm2* expression in this projection (Fig 1B and Fig. 5A).

**Figure 5:**
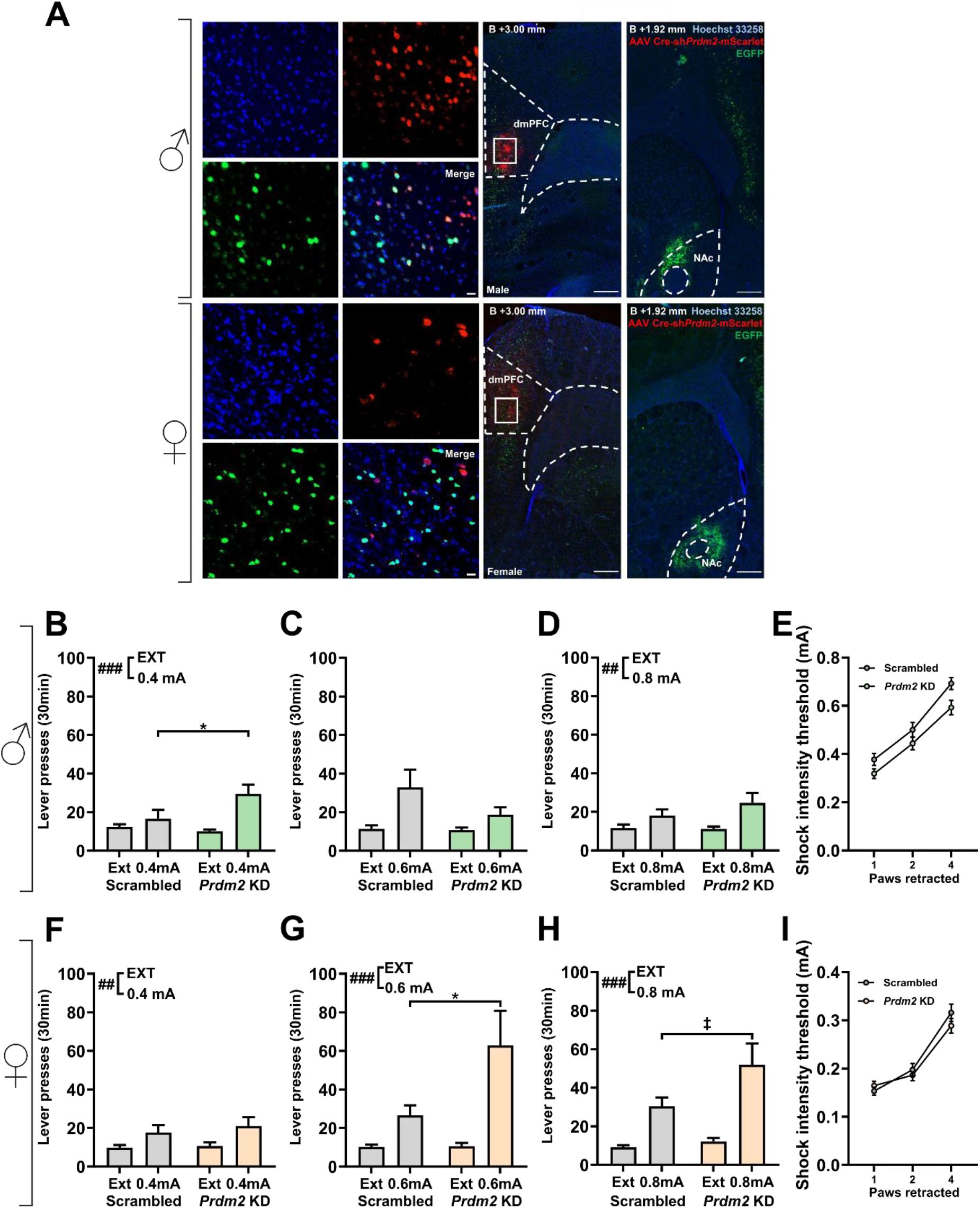
*Prdm2* knockdown in dmPFC→NAc projection neurons regulates stress-induced reinstatement of alcohol seeking in male and female rats. (A) Representative images showing viral targeting of the dorsomedial prefrontal cortex (dmPFC) and the nucleus accumbens (NAc). Left: 20x magnification image of AAV5-hSyn-mScarlet-shPrdm2-DIO expression (mScarlet, red) and retrogradely transported AAV encoding Cre (EGFP, green) with nuclear staining (Hoechst 33258, blue). Scale bar: 50 µm Right: schematic showing viral expression restricted to the dmPFC and the NAc. Scale bar: 400 µm. (B–D) Stress-induced reinstatement of alcohol seeking in male rats following intermittent footshock at 0.4 mA (B), 0.6 mA (C), and 0.8 mA (D). Stress exposure significantly increased lever pressing compared with extinction conditions across shock intensities. *Prdm2* knockdown in dmPFC→NAc neurons significantly enhanced reinstatement responding at 0.4 mA, but not at 0.6 or 0.8 mA. (E) Footshock sensitivity thresholds in male rats measured as the current intensity required to elicit increasing paw withdrawal responses (1, 2, or 4 paws retracted). No differences were observed between scrambled and *Prdm2* KD groups. (F–H) Stress-induced reinstatement of alcohol seeking in female rats tested at 0.4 mA (F), 0.6 mA (G), and 0.8 mA (H). *Prdm2* knockdown in dmPFC→NAc neurons significantly increased reinstatement responding at 0.6 mA and showed a trend toward significance at 0.8 mA, but not at 0.4 mA. (I) Footshock sensitivity thresholds in female rats. *Prdm2* knockdown did not alter pain sensitivity to footshock. Bars represent mean ± SEM lever presses during the 30-min session. ‡*p* = 0.058 (compared to scrambled control); \**p* < 0.01 (compared to scrambled control); ## *p* < 0.01, ### *p* < 0.001 main effect of stress compared with extinction.

All rats underwent footshock-induced reinstatement testing at 0.4, 0.6, and 0.8 mA shock intensities (males: Fig. 5B-E; females: Fig. 5F-I). In males, stress exposure significantly reinstated alcohol seeking at 0.4 mA and 0.8 mA shock intensities. Importantly, *Prdm2* KD male rats show significantly greater stress-induced reinstatement at 0.4 mA compared with scrambled controls. At 0.4 mA, repeated-measures ANOVA revealed a significant main effect of stress (F_(1,35)_=13.10, p=0.0009), and a significant Stress × Group interaction (F_(1,35)_=5.24, p=0.028), with no main effect of group (F_(1,35)_=2.19, p=0.15; Fig. 5B). Post hoc comparisons indicated that dmPFC→NAc *Prdm2* KD rats showed greater reinstatement responding compared with scrambled controls (P=0.001). At 0.6 mA, because homogeneity of variance assumption was violated (Levene’s test: p=0.00028), data were analyzed using an ART ANOVA. This analysis showed no significant main effect of group (F_(1,36)_=1.45, p=0.24), a trend for a main effect of stress (F_(1,36)_=3.36, p=0.075), and no Group × Stress interaction (F_(1,36)_=1.82, p=0.19; Fig. 5C). At 0.8 mA, repeated-measures ANOVA showed a significant main effect of stress (F_(1,37)_=11.28, p=0.0018), but no main effect of group (F_(1,37)_=0.74, p=0.40) and no Stress × Group interaction (F_(1,37)_=1.40, p=0.24; Fig. 5D). *Prdm2* KD in dmPFC-NAc projecting neurons did not change pain sensitivity to footshocks in males (repeated-measures ANOVA: no main effect of Group, F_(1,21)_=3.15,p=0.09, and no Group × Intensity interaction, F_(2,42)_=0.49, p=0.62; Fig. 5E), indicating that the differences in reinstatement between groups does not result from altered pain thresholds.

Similarly, *Prdm2* KD females exhibited significantly greater stress-induced reinstatement at 0.6 mA compared with scrambled controls. At 0.4 mA, repeated-measures ANOVA revealed a significant main effect of stress (F_(1,40)_=10.82, p=0.0021), but no main effect of group (F_(1,40)_=0.36, p=0.55) and no Stress × Group interaction (F_(1,40)_=0.20, p=0.66; Fig. 5F). At 0.6 mA, because variance homogeneity assumptions were violated (Levene’s test: p=0.00077), data were analyzed using ART ANOVA, which showed a significant main effect of group (F_(1,42)_=5.40, p=0.025), a significant main effect of stress (F_(1,42)_=18.86, p=8.7×10^−5^), and a significant Group × Stress interaction (F_(1,42)_=6.62, p=0.0137; Fig. 5G). Post hoc comparisons indicated that dmPFC→NAc *Prdm2* KD females exhibited greater reinstatement responding compared with scrambled controls (P=0.02). At 0.8 mA, ART ANOVA showed a significant main effect of stress (F_(1,41)_=63.47, p=7.37×10^−10^), but no main effect of group (F_(1,41)_=3.78, p=0.058) and no Group × Stress interaction (F_(1,41)_=2.88, p=0.097; Fig. 5H). *Prdm2* KD in dmPFC→NAc projecting neurons did not change pain sensitivity to footshocks in females (repeated-measures ANOVA: no main effect of Group, F_(1,9)_=0.33, p=0.58, and no Group × Interaction, F_(3,27)_=1.03, p=0.39; Fig. 5I), indicating that the differences in reinstatement between groups does not result from altered pain thresholds.

Together, these findings indicate that reducing *Prdm2* expression in dmPFC neurons projecting to the NAc enhances stress-induced reinstatement of alcohol seeking in a circuit- and intensity-dependent manner, supporting a role for the dmPFC→NAc pathway in mediating PRDM2-dependent regulation of stress-induced reinstatement.

## DISCUSSION

Our findings identify *PRDM2* as a regulator of stress-induced reinstatement of alcohol seeking and provide insight into circuit mechanisms linking prefrontal dysfunction to alcohol use disorder. Consistent with our previous findings in post-dependent rats (Barbier et al., 2017), we found that *PRDM2* expression is similarly decreased in the PFC of individuals with AUD, suggesting that reduced *PRDM2* expression may represent a conserved molecular feature of alcohol dependence across species. Building on this cross-species observation, our preclinical experiments provide mechanistic insight into how reduced PRDM2 function contributes to relapse-like behavior. *Prdm2* KD in the dmPFC enhanced stress-induced reinstatement of alcohol seeking in male rats, implicating the dmPFC→NAc circuitry in the regulation of relapse-like behavior. Importantly, we show that this effect also occurs in female rats, indicating that *PRDM2* regulates stress-induced reinstatement through mechanisms that are conserved across sexes. Together, these findings suggest that alcohol-induced reductions in *Prdm2* expression may contribute to maladaptive transcriptional states within the PFC that increase stress-induced reinstatement.

The reduction of *PRDM2* expression in the PFC of individuals with AUD converges with findings from rodent models and supports the translational relevance of these preclinical observations. Consistent with the central role of the PFC in executive control and regulation of reward-related behaviors (Goldstein and Volkow, 2011; Miller and Cohen, 2001), reduced *PRDM2* expression may disrupt PFC circuits involved in stress and motivation, thereby contributing to relapse. These cross-species findings should be interpreted in light of differences in anatomical resolution between the datasets. In rodents, our manipulations targeted the dmPFC, encompassing the prelimbic cortex implicated in relapse-related behavior. In contrast, *PRDM2* expression in humans was assessed in PFC tissue spanning Brodmann areas 9, 10, and 46. The rodent mPFC is generally regarded as functionally homologous to the dorsolateral prefrontal cortex in humans and nonhuman primates (Farovik et al., 2008), suggesting that the human and rodent findings engage related prefrontal systems despite anatomical differences. We found that females showed significantly lower levels of *PRDM2* expression in the PFC compared to males. Although stress and negative affect contribute to all phases of AUD in both sexes, their impact is higher in women (Verplaetse et al., 2018) and this might be associated with reduced *PRDM2* expression.

While activity of the dmPFC has been shown to be involved in stress-induced reinstatement of alcohol seeking, the specific prefrontal output pathways mediating this behavior remain poorly defined. The dmPFC projection to the NAc is a strong candidate circuit, which has been shown to regulate stress-induced reinstatement of cocaine seeking (McFarland et al., 2004). However, whether this projection similarly contributes to stress-induced alcohol seeking has not been directly tested. Our projection-specific approach shows that reducing *Prdm2* expression in dmPFC neurons projecting to the NAc enhances stress-induced reinstatement of alcohol seeking, identifying the dmPFC→NAc pathway as a key circuit node contributing to alcohol relapse-like behavior. These findings therefore extend the involvement of this relapse circuit from psychostimulants to alcohol and suggest that PRDM2-dependent transcriptional regulation within dmPFC→NAc neurons may modulate the activity of this projection to promote stress-induced reinstatement of alcohol seeking.

Notably, the behavioral effects of *Prdm2* knockdown were dependent on stress intensity. When *Prdm2* expression was reduced in the dmPFC, enhanced reinstatement was most prominent at the intermediate shock intensity (0.6 mA) in both male and female rats. One interpretation is that reduced *PRDM2* expression enhances the engagement of relapse-related circuits at stress intensities that effectively recruit, but do not saturate, reinstatement behavior. At lower shock intensities (0.4 mA), the stressor may be insufficient to robustly recruit reinstatement behavior, whereas at higher intensities (0.8 mA), stress may already produce near-maximal reinstatement, limiting the ability to detect further enhancement. In our previous study, dmPFC *Prdm2* knockdown enhanced stress-induced reinstatement at both 0.4 and 0.6 mA shock intensities (Barbier et al., 2017). This discrepancy may reflect methodological differences between studies, including the use of a saccharin-fading alcohol self-administration procedure, a FR1 schedule of reinforcement rather than FR2, and a different viral vector system. Interestingly, projection-specific knockdown of *Prdm2* in dmPFC→NAc neurons enhanced reinstatement at lower stress intensity in males (0.4 mA), whereas in females the effect was observed at the intermediate intensity (0.6 mA). These findings suggest that PRDM2 modulates the recruitment of prefrontal-striatal circuits that promote relapse-like behavior in a stress intensity-dependent manner.

At the cellular level, we previously demonstrated that PRDM2 regulates transcriptional signatures linked to synaptic plasticity (Barchiesi et al., 2022). Reduced *PRDM2* expression may therefore alter excitatory signaling within cortico-striatal circuits that govern stress-induced reinstatement. Although the precise synaptic mechanisms remain to be determined, PRDM2-dependent transcriptional regulation could influence activity within the dmPFC-NAc pathway by altering the expression of genes involved in synaptic transmission.

Together, the present findings highlight PRDM2 as a potential molecular link between chronic alcohol exposure and dysregulation of relapse circuits. Epigenetic mechanisms are increasingly recognized as key regulators of long-lasting neural adaptations in addiction (Domi et al., 2025; Robinson and Nestler, 2011), and targeting epigenetic regulators may provide a strategy for restoring transcriptional control within maladaptive circuits. Our results suggest that reduced *PRDM2* expression in the PFC may increase the sensitivity of cortico-striatal relapse circuits to stress, a major trigger of relapse in individuals with AUD. Importantly, the observation that PRDM2 regulates stress-induced reinstatement in both male and female animals indicates that this mechanism is conserved across sexes, highlighting its potential relevance for developing treatments that may be effective in both men and women. By identifying PRDM2 as a regulator of stress-induced relapse-like behavior and linking its function to a defined prefrontal projection, this study provides a framework for understanding how alcohol-induced epigenetic adaptations in the PFC may promote relapse. Together with our prior findings implicating *PRDM2* in fear regulation (Barchiesi et al., 2022), these results support a broader role for *PRDM2*-dependent epigenetic mechanisms in alcohol- and stress-related disorders across sexes and potentially across species.

## ACKNOWLEDGEMENT

We thank Prof. Markus Heilig for his valuable scientific comments and Lovisa Holms for her technical support. We also thank Dr. Vesa Loitto at the core facility at the Faculty of Medicine and Health Sciences, Linköping University for microscopy technical assistance. This study was supported by Council Swedish Research Council 2023-02315.

## CONTRIBUTION

EB, KC, and NM conceptualized and designed the studies. EB designed and supervised experiments and interpreted results. EB, KC, NM, and LX performed surgeries. KC and NM performed the behavioral experiments. KC, NM, and LX performed the molecular analysis. KC, NM, STE, TK, SV and AC contributed to the data collection. EB, ED, KC and NM drafted and revised the manuscript. All authors have read and approved the manuscript.

## FINANTIAL DISCLOSURE

The authors has no financial disclosure.

