## Supplementary information for "Reduced Prefrontal PRDM2 Promotes Stress-Induced Alcohol Reinstatement Across Sexes through a dmPFC-Nucleus Accumbens Pathway"

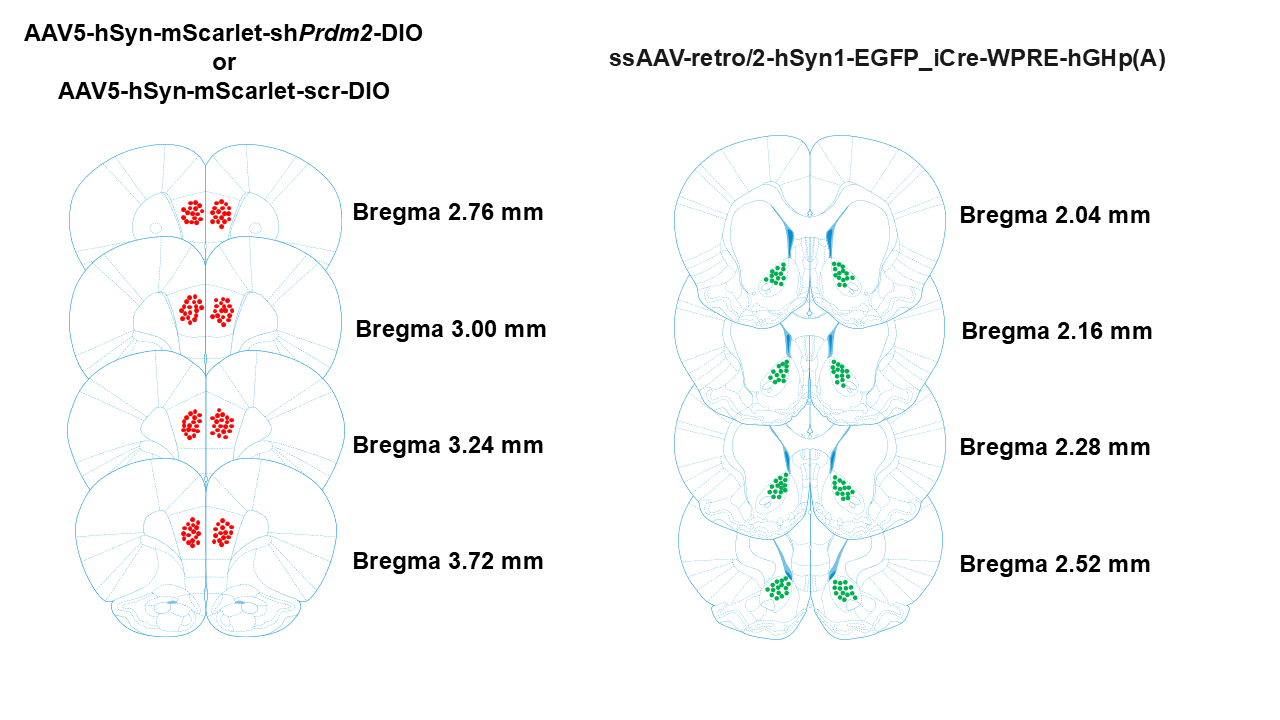

**Fig. S1.** Representative images depicting targeted coordinates (related to bregma in mm) in dmPFC (red) and NAc (green).

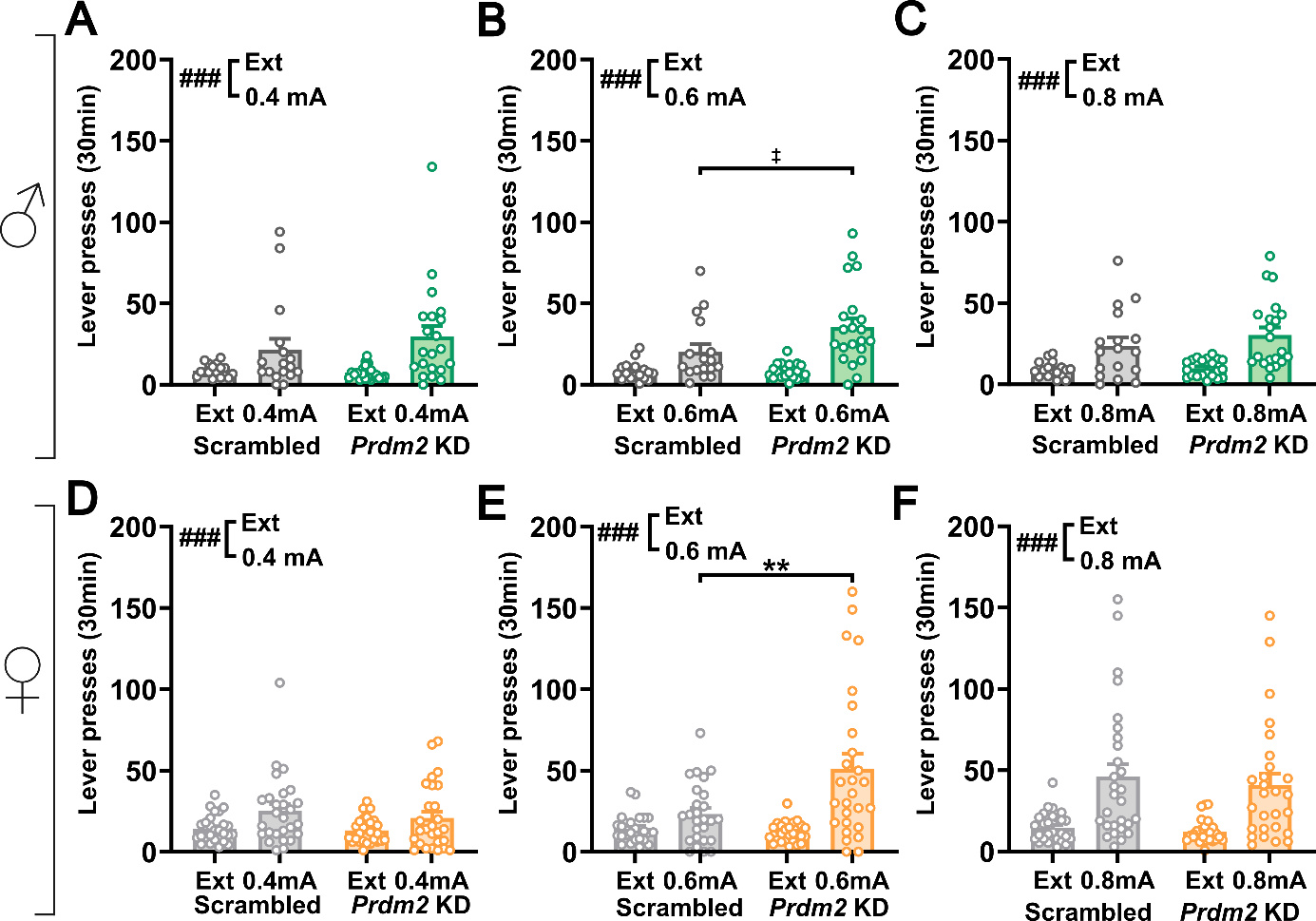

**Fig. S2. Knockdown of *Prdm2* in the dmPFC enhances stress-induced reinstatement of alcohol seeking at intermediate shock intensity in both male and female rats.** (A–C) Individual values in stress-induced reinstatement of alcohol seeking in male rats following intermittent footshock at 0.4 mA (A), 0.6 mA (B), and 0.8 mA (C). Stress exposure increased lever pressing relative to extinction conditions across shock intensities, with *Prdm2* KD significantly enhancing reinstatement responding at 0.6 mA. (D-F) Individual values in stress-induced reinstatement of alcohol seeking in female rats tested at 0.4 mA (D), 0.6 mA (E), and 0.8 mA (F). Similar to males, *Prdm2* KD significantly increased reinstatement responding at 0.6 mA but not at the lower or higher shock intensities. Bars represent mean ± SEM lever presses during the 30-min session. ‡ *p* = 0.05 (compared to scrambled control), ** *p* < 0.01 (compared to scrambled control); ### *p* < 0.001 main effect of stress compared with extinction.

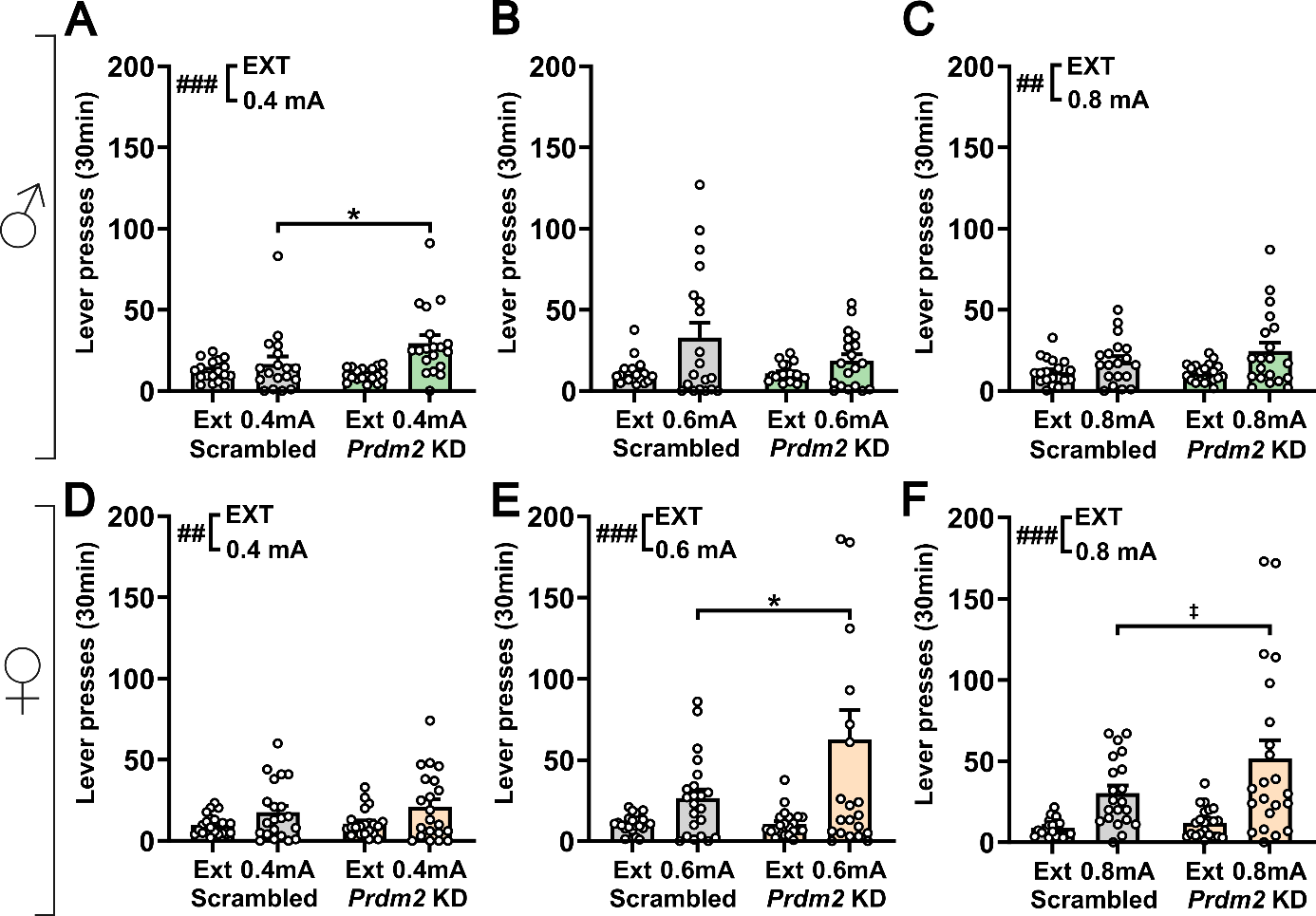

**Fig. S3. PRDM2 in dmPFC_→_NAc projection neurons regulates stress-induced reinstatement of alcohol seeking in male and female rats.** (A–C) Individual values in stress-induced reinstatement of alcohol seeking in male rats following intermittent footshock at 0.4 mA (A), 0.6 mA (B), and 0.8 mA (C). Stress exposure significantly increased lever pressing compared with extinction conditions across shock intensities. *Prdm2* knockdown in dmPFC_→_NAc neurons significantly enhanced reinstatement responding at 0.4 mA, but not at 0.6 or 0.8 mA. (D-F) Individual values in stress-induced reinstatement of alcohol seeking in female rats tested at 0.4 mA (D), 0.6 mA (E), and 0.8 mA (F). *Prdm2* knockdown in dmPFC_→_NAc neurons significantly increased reinstatement responding at 0.6 mA and showed a trend toward significance at 0.8 mA, but not at 0.4 mA. Bars represent mean ± SEM lever presses during the 30-min session. ‡ *p* = 0.058; **p* < 0.01; ## *p* < 0.01, ### *p* < 0.001 main effect of stress compared with extinction.
